# Field-based species identification in eukaryotes using real-time nanopore sequencing

**DOI:** 10.1101/107656

**Authors:** Joe Parker, Andrew J. Helmstetter, Dion Devey, Alexander S.T. Papadopulos

**Affiliations:** Jodrell Laboratory, Royal Botanic Gardens, Kew, Richmond, Surrey UK. TW9 3A

**Keywords:** Nanopore, MinlON, onsite DNA sequencing, phylogenomics

## Abstract

Advances in DNA sequencing and informatics have revolutionised biology over the past four decades, but technological limitations have left many applications unexplored^1,2^. Recently, portable, real-time, nanopore sequencing (RTnS) has become available. This offers opportunities to rapidly collect and analyse genomic data anywhere^3–5^. However, the generation of datasets from large, complex genomes has been constrained to laboratories^6,7^. The portability and long DNA sequences of RTnS offer great potential for field-based species identification, but the feasibility and accuracy of these technologies for this purpose have not been assessed. Here, we show that a field-based RTnS analysis of closely-related plant species (*Arabidopsis spp*.)^8^ has many advantages over laboratory-based high-throughput sequencing (HTS) methods for species level identification-by-sequencing and *de novo* phylogenomics. Samples were collected and sequenced in a single day by RTnS using a portable, *“al fresco”* laboratory. Our analyses demonstrate that correctly identifying unknown reads from matches to a reference database with RTnS reads enables rapid and confident species identification. Individually annotated RTnS reads can be used to infer the evolutionary relationships of *A. thaliana*. Furthermore, hybrid genome assembly with RTnS and HTS reads substantially improved upon a genome assembled from HTS reads alone. Field-based RTnS makes real-time, rapid specimen identification and genome wide analyses possible. These technological advances are set to revolutionise research in the biological sciences^9^ and have broad implications for conservation, taxonomy, border agencies and citizen science.

---

DNA sequencing used to be a slow undertaking, but the past decade has seen an explosion in HTS methods^2,10^. DNA barcoding (i.e., the use of a few, short DNA sequences to identify organisms) has benefited from this sequencing revolution^11,12^, but has never become fully portable. Samples must be returned to a laboratory for testing and the discrimination of closely related species using few genes can be problematic due to evolutionary phenomena (e.g. lineage sorting, shared polymorphism and hybridisation)^10^. While typical barcoding approaches have been effective for generic level identification, accuracy is much more limited at the species level^11,13^ and concerns remain^14^. Species delimitation using limited sequencing information has also been problematic and is thought to heavily underestimate species diversity^11,15^. Consequently, increasingly elaborate analytical methods have been spawned to mitigate the inherent limitations of short sequences^13,16^. The Oxford Nanopore Technologies^®^ MinION^®^ is one of a new generation of RTnS DNA sequencers that is small enough to be portable for fieldwork and produces data within minutes^17,18^. These properties suggest species identification could be conducted using genome scale data generated at the point of sample collection. Furthermore, the large number of long reads generated^17^ may provide more accurate species-level identification than current approaches. This application offers great potential for conservation, environmental biology, evolutionary biology and combating wildlife crime, however, this potentially exciting combination of methods has not yet been tested in the field for eukaryotes.

Our experiment was designed to determine whether DNA reads produced entirely in the field could accurately identify and distinguish samples from closely-related species (*A. thaliana* (L.) Heynh. and *A. lyrata* (L.) O’Kane & Al-Shehbaz). Recent analyses have shown that gene flow has been common and shared polymorphisms are abundant between the morphologically distinct species in *Arabidopsis.* Indeed, the two study species share >20,000 synonymous SNPs^8^, making this a good stress test of genome scale RTnS sequencing for species discrimination.

The first goal was to extract and sequence shotgun genomic data from higher plant species in the field using RTnS technology in sufficient quantity for downstream analyses within hours of the collection of plant tissue (Extended Figure 1). On consecutive days, tissue was collected from three specimens each of *A. thaliana* and *A. lyrata subsp. petraea* (Figs. 1b,c) in Snowdonia National Park, and prepared, sequenced and analysed outdoors in the Croesor Valley (Fig. 1a). Only basic laboratory equipment was used for DNA extraction and MinlON sequencing-library preparation; we did not use a PCR machine (Fig. 1d; Extended Data Table 1). One specimen of each species was sequenced with both R7.3 and R9 MinlON chemistries. For *A. thaliana*, the RTnS experiment generated 97k reads with a total yield of 204.6Mbp over fewer than 16h of sequencing (see Extended Data Table 2). Data generation was slower for *A. lyrata*, over ~90h sequencing (including three days of sequencing at RBG Kew following a 16h drive), 26k reads were generated with a total yield of 62.2Mbp. At the time, a limited implementation of local basecalling was available for the R7.3 data only. Of 1,813 locally basecalled reads, 281 had successful BLAST matches to the reference databases with a correct to incorrect species ID ratio of 223:30. The same samples were subsequently sequenced using HTS short read technology (Illumina MiSeq™, paired-end, 300bp; Supplementary note 1). Mapping reads to available reference genomes for the *A. thaliana* (TAIR10 release^19^) and two *A. lyrata* assemblies^20,21^ indicates approximate RTnS coverage of 2.0x, 0.3x, and 0.3x for *A. thaliana, A. lyrata*, and *A. lyrata ssp. petraea*, respectively; and 19.5x, 11.9x and 12.0x respectively for HTS reads (Extended Data Tables 2 and 3, Supplementary note 2). These results demonstrate that the entire process (from sample collection thorough to genome scale sequencing) is now feasible for eukaryotic species within a few hours in field conditions. As the technology develops, run yields are expected to improve and implementation of sample indexing will allow many samples to be run on a single flow cell.

**Figure 1.**
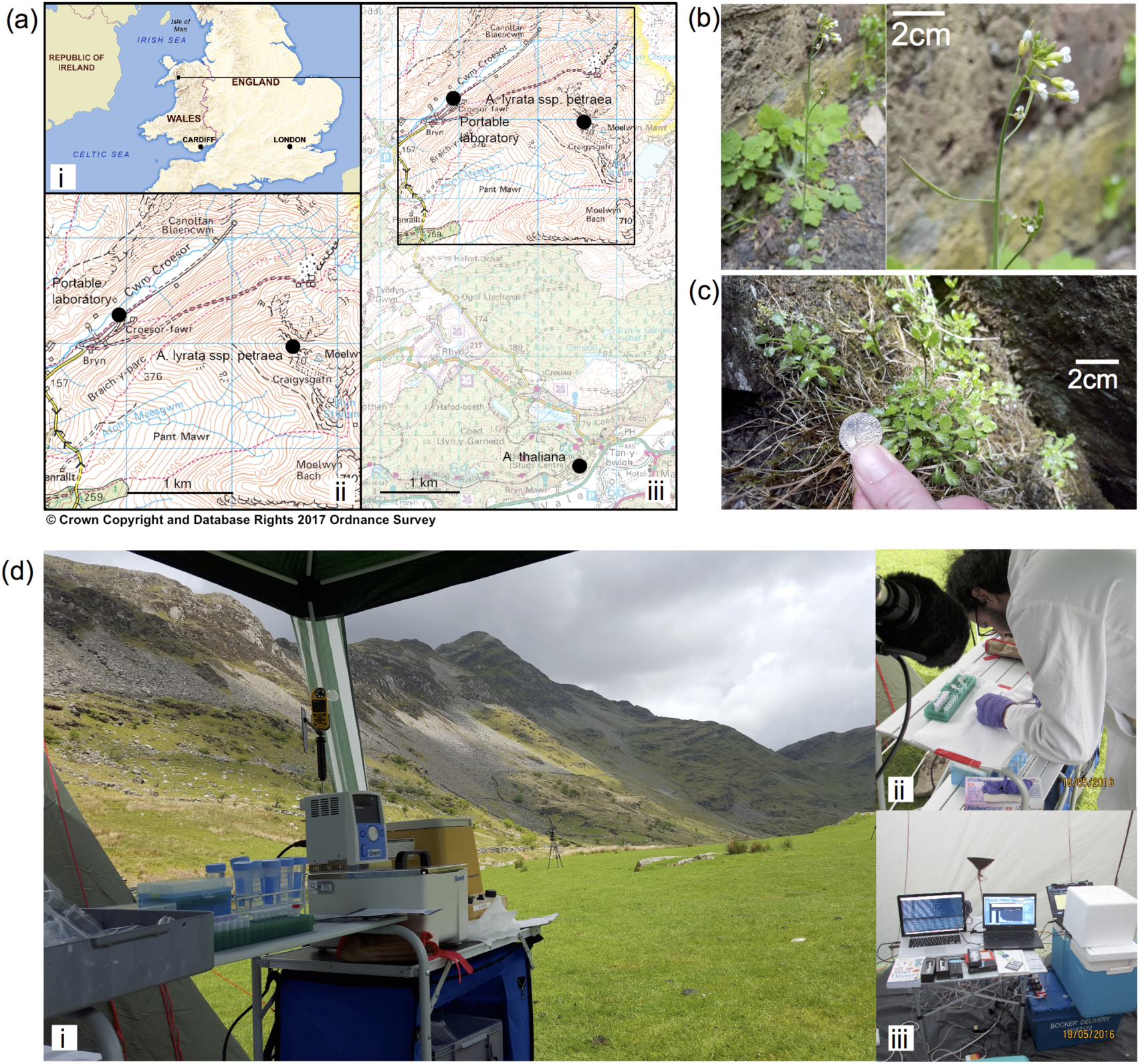
Logistics and scope of field-based sequencing. **a,** Location of sample collection and extraction, sequencing and analyses in the Snowdonia National Park, Wales. **b,** *Arabidopsis thaliana.* **c,** *A. lyrata ssp. petraea.* **d,** The portable field laboratory used for the research. Ambient temperatures varied between 7-16°C with peak humidity >80%. A portable generator was used to supply electrical power.

As expected given the developmental stages of the technologies, the quality and yield of field sequenced RTnS data was lower than the HTS data (Extended Data Tables 2 and 4). *Arabidopsis thaliana* RTnS reads could be aligned to approx. 50% of the reference genome (53Mbp) with an average error rate of 20.9%. Indels and mismatches were present in similar proportions. The *A. lyrata* RTnS data were more problematic with significantly poorer mapping to the two *A. lyrata* assemblies, whereas, the HTS data performed relatively well. For the limited number of alignable RTnS reads, error rates were slightly higher than for *A. thaliana* (22.5% and 23.5%). The poorer RTnS results for *A. lyrata* may be a consequence of temperature-related reagent degradation in the field or due to unknown contaminants in the DNA extraction that inhibited library preparation and/or RTnS sequencing. Despite the smaller yield and lower accuracy of the RTnS compared to HTS data, the RTnS reads were up to four orders of magnitude longer than the HTS reads and we predicted they would be useful for species identification, hybrid genome assembly and phylogenomics.

To explore the utility of these data for species identification, the statistical performance of field-sequenced (RTnS) and lab-sequenced (HTS) read data was assessed. Datasets for each species were compared to two databases via BLASTN, retaining single best-hits: one database contained the *A. thaliana* reference genome and the second was composed of the two draft *A. lyrata* genomes combined. Reads which matched a single database were counted as positive matches for that species. The majority of matching reads hit both databases, which is expected given the close evolutionary relationships of the species. In these cases, positive identifications were determined based on four metrics; a) the longest alignment length, b) the highest % sequence identities and c) the largest number of sequence identities d) the lowest E-value (Extended Data Table 4). Test statistics for each of these metrics were calculated as the difference of scores (length, *%* identities, or E-value) between ‘correct’ and ‘incorrect’ database matches. The performance of these difference statistics for binary classification was assessed by investigating the true and false positive rates (by reference to the known sample species) across a range of threshold difference values (Fig. 2a-d and Extended Data Figs. 2–4). For both short-and long-read data at thresholds greater than 100bp, the differences in total alignment lengths (ΔL_T_) or number of identities (ΔL_I_) are superior to e-value or % identity biases (Figs. 2a-d). Furthermore, at larger thresholds (i.e., more conservative tests), RTnS reads retained more accuracy in true-and false-positive discrimination than HTS data. This proves that whole genome shotgun RTnS is a powerful method for species identification. We posit that the extremely long length of the observed ‘true positive’ alignments compared with an inherent length ceiling on false-positive alignments in a typical BLASTn search is largely responsible for this property.

**Figure 2.**
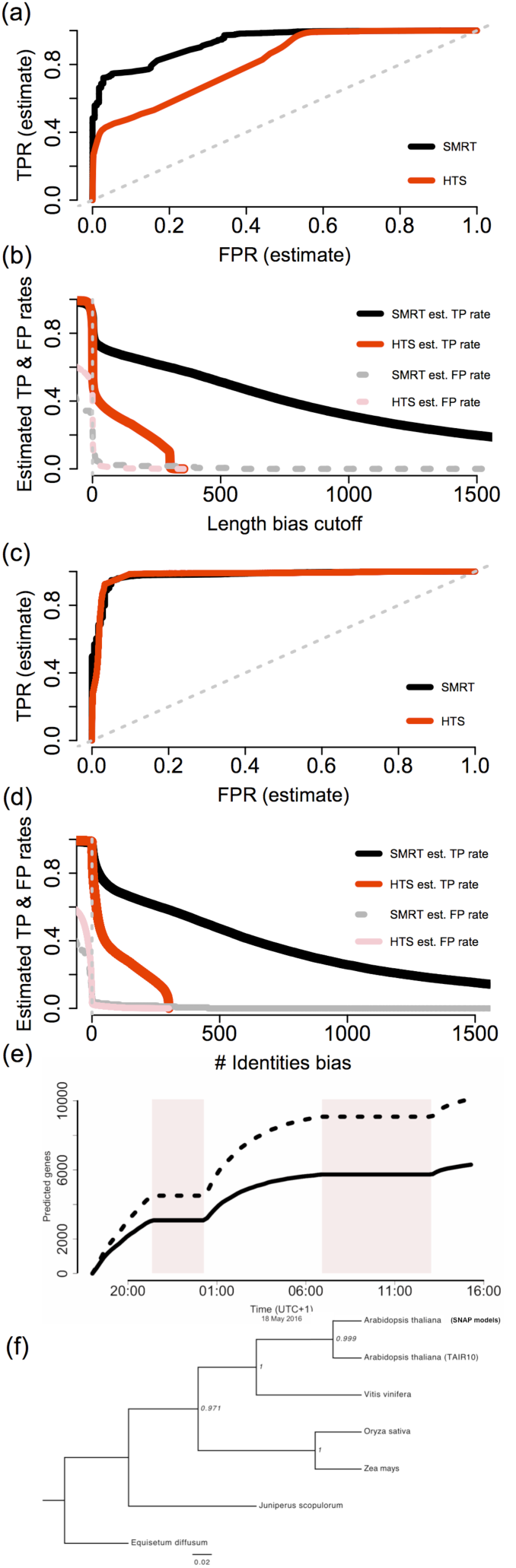
Sample identification and phylogenomics using field-sequenced RTnS data. **a-d** Orthogonal species identification using BLASTN difference statistics: HTS data (red) and RTnS (black) matched to reference databases via BLASTN. **a, c** Receiver operating characteristic (ROC; estimated false-positive rate vs. estimated true positive rate) and **b, d** estimated true-(solid lines) and false-positive (dashed lines) rates. **a, b** ΔL_T_ statistic; **c, d,** ΔL_I_ statistic. **e,** Accumulation curves for *ab initio* gene models predicted directly from individual *A. thaliana* reads over time. Count of unique TAIR10 genes (solid line) and total number of gene models (dashed line). Shaded boxes represent periods where the MinION devices were halted while the laboratory was dismantled and moved. **f,** phylogenetic tree inferred under the multispecies coalescent from RTnS reads.

To evaluate the speed with which species identification can be carried out, we performed *post hoc* analyses by subsampling the RTnS *A. thaliana* dataset. This simulated the rate of improvement in species assignment confidence over a short RTnS run. We classified hits among the subsampled reads based on (i) ΔL_I_ over a range of threshold values (ii) mean ΔL_I_ and (iii) aggregate ΔL_I_ (Fig. 3). This demonstrates that a high degree of confidence can be assigned to species identifications over the timescales needed to generate this much data (i.e., < one hour) and that variation in the accuracy of identifications quickly stabilises above 1000 reads. Aggregate ΔL_I_ values rapidly exclude zero (no signal) or negative (incorrect assignment) values, making this simple and rapidly-calculated statistic particularly useful for species identification. In a multispecies context, the slopes of several such log-accumulation curves could be readily compared, for example (see Supplementary Discussion).

**Figure 3.**
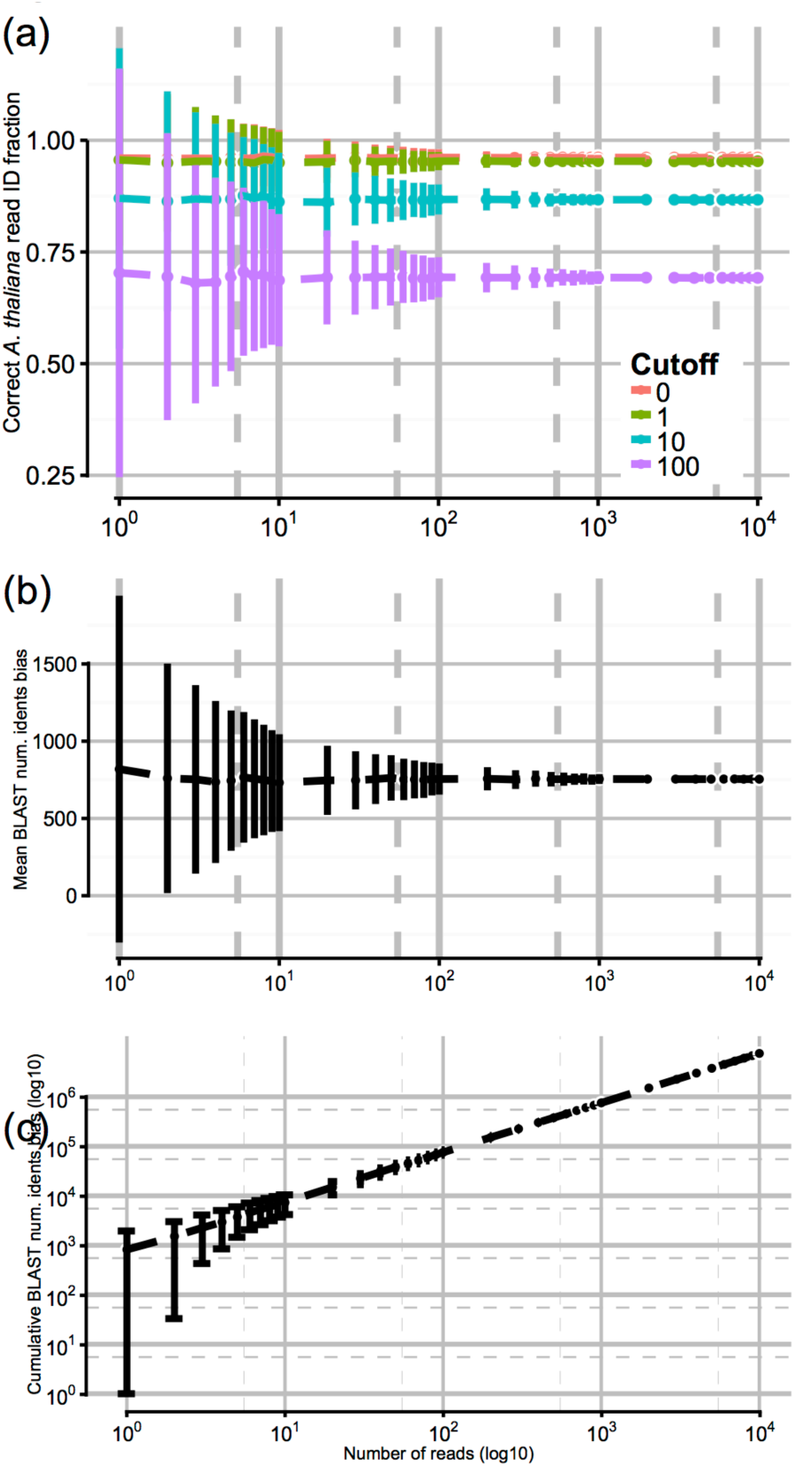
Simulated accumulation curves for rapid species identification by DNA sequencing. 34k pairwise BLASTN hits of *A. thaliana* RTnS reads were subsampled without replacement to simulate an incremental accumulation of data (10^4^ reads; 10^3^ replicates). For each read the total identities bias (ΔL_I_) is the number of identities with the *A. thaliana* reference minus the number of identities with the *A. lyrata* reference. **a)** the proportion of *A. thaliana* reads correctly identified on a per-read basis, classified as *A. thaliana* where ΔL_I_ > threshold cutoff (0, 1, 10 or 100). **b)** Mean ΔL_I_ in the simulated dataset rapidly stabilises on the population mean (+754bp, e.g. an average matching read alignment to *A. thaliana* is 754bp longer than to *A. lyrata*). **c)** Cumulative aggregate ΔL_I_; negative or zero ΔL_I_ can rapidly be excluded. Typical data throughput rates exceed 10^4^ reads per hour of sequencing.

Assembly of large and complicated eukaryotic genomes with RTnS data alone would require a greater volume of data than available here^7, 22–24^. Field extracted samples are unlikely to be of similar purity to those obtained with more sophisticated laboratory-based methods, leading to lower yields. As expected, *de novo* assembly of our RTnS data performed poorly, likely due to insufficient coverage. However, these data do have potential for hybrid genome assembly approaches. We assembled the HTS data *de novo* using ABYSS^25^ and produced a hybrid assembly with both RTnS and HTS datasets using HybridSPAdes^26^. The hybrid assembly was an improvement over the HTS-only assembly (see Extended Data Table 5) with fewer contigs, a total assembly length closer to the reference (119.0Mbp), N50 and longest contig statistics both increasing substantially and estimated completeness (CEGMA^27^) of coding loci increased to ~99%. These results suggest that relatively small quantities of long and short reads can produce useful genome assemblies when analysed together, an important secondary benefit of field–sequenced data. The length of typical RTnS reads is similar to that of genomic coding sequences (1-10kb)^17^. This raises the possibility of extracting useful phylogenetic signal from such data, despite the relatively high error rates of individual reads. We annotated individual raw *A. thaliana* reads directly, without genome assembly, which recovered over 2,000 coding loci from the data sequenced in the first three hours (Fig. 2e). These predicted gene sequences were combined with a published dataset spanning 852 orthologous, single-copy genes^28^, downsampled to 6 representative taxa. Of our gene models, 207 were present in the Wickett *et al*.^28^ dataset and the best 56 matches were used for phylogenomic analysis (see Supplementary Methods for details). The resulting phylogenetic trees (Fig. 2f and Extended Data Fig. 5) are consistent with the established intergeneric relationships^28^. Although the taxonomic scale used here for phylogenomics is coarse it highlights an additional benefit to rapid, in-the-field sequencing for evolutionary research.

This experiment is the first to demonstrate field-based sequencing of higher plant species. When directly compared to lab-based HTS, our experiment highlights key discriminatory metrics for highly accurate species identifications using portable RTnS sequencing. Few approaches can boast this level of discriminatory power and none of these have the same degree of portability^10,11^. The data produced for identification is also useful for genome assembly. Entire coding sequences can be recovered from single reads and incorporated into evolutionary analyses. Clearly, data generated with the goal of accurate species identification has much broader usefulness for genomic and evolutionary research. Few technical barriers remain to prevent the adoption of portable RTnS by non-specialists, or even keen amateurs and schoolchildren. As these tools mature, and the number of users expands, portable RTnS sequencing can revolutionise the way in which researchers and practitioners can approach ecological, evolutionary and conservation questions.

## Methods summary

Genomic DNA was extracted from two plant specimens and sequenced on Oxford Nanopore MinION devices according to manufacturers’ recommendations in a portable outdoor laboratory. Offline basecalling software and local BLAST (v2.2.31) were used to identify individual reads on-site. Short reads were sequenced in the laboratory from the same extracted DNA using an Illumina MiSeq. Local BLAST was used to identify reads from all four datasets (2 field × 2 species) by comparison to available published reference genomes. Gene models were predicted directly from individual DNA reads using SNAP (v2006-07-28), matched to existing phylogenomic datasets and used to infer plant phylogenies using MUSCLE (v3.8.31) and RAxML (v7.2.8). *de novo* genome assemblies were performed using Abyss (v1.5.2) and Hybrid-SPAdes (v3.5.1) with completeness assessed with QUAST (v4.0) and CEGMA. R (v3.1.3) was used to perform statistical analyses. Additional details are given in the Supplementary Methods.

## Acknowledgements

This work was funded by a Pilot Study Grant to JDP and a Howard Lloyd Davies legacy grant to ASTP. JDP was also supported by funding from the Calleva Foundation Phylogenomic Research Programme and the Sackler Trust. The authors also thank The Botanical Society of Britain & Ireland. Natural Resources Wales, Tim Wilkinson, Robyn Cowan, and Patricia and David Brandwood for assistance.

## Author contributions

ASTP and JDP conceived the study and obtained funding. ASTP, DD and JDP designed and conducted fieldwork. ASTP designed and conducted field-based labwork with input from JDP, AH and DD. AH conducted lab-based sequencing. JDP conducted bioinformatics and phylogenomic analyses with contributions from AH. ASTP and JDP prepared the manuscript with contributions from DD and AH.

## Author information

Basecalled read data for Illumina and Oxford Nanopore sequencing runs are available via the EBI ENA at XXXXX. The authors declare competing financial interests: Oxford Nanopore Technologies contributed MinION sequencing reagents and flowcells for this research. JDP and ASTP received travel remuneration and free tickets to present an early version of this work at a conference (London Calling 2016). Correspondence and requests for materials should be addressed to ASTP and JDP.

**Supplementary Information** is linked to the online version of the paper.

## Extended Methods

### Study site and sample collection

On consecutive days, tissue was collected from three specimens each of *A. thaliana* and *A. lyrata subsp. petraea* in Snowdonia National Park and sequenced and analysed in a tent. *A. lyrata* was collected from the summit of Moelwyn Mawr (52.985168° N, 4.003754° W; OL17 65554500; SH6558244971) and *Arabidopsis thaliana* was collected at Plâs Tan-y-Bwlch (52.945976° N 4.002730° W; OL18 65604060; SH6552940610). Representative voucher specimens of each species are deposited at RBG, Kew. DNA extractions, library preparation and DNA sequencing with the MinlON technology were all conducted using portable laboratory equipment in the Croesor valley on the lower slopes of Moelwyn Mawr immediately following sample collection (52.987463°N 4.028517° W; OL17 63904530; SH6392745273). Laboratory reagents were stored in passively-cooled polystyrene boxes with internal temperatures monitored using an Arduino Uno. Only basic laboratory equipment was used (including two MinlON sequencers and three laptops; see Extended Data Table 1).

### DNA extraction

The standard Qiagen DNeasy plant mini prep kit was used to extract genomic DNA from *Arabidopsis spp.* with the exception that the two batches were pooled at the DNeasy mini spin column step to maximise the DNA yield.

### DNA library preparation and sequencing

An R7.3 and R9 1D MinlON library preparation were performed for each species according to the manufacturer’s instructions using a developer access programme version of the commercially available Nanopore RAD-001 library kit (Oxford Nanopore Technologies). No PCR machine was used. Lambda phage DNA was added to *A. thaliana* R9 library for quality control. For *A. thaliana*, the MinION experiment generated 96,845 1D reads with a total yield of 204.6Mbp over fewer than 16h of sequencing. Data generation was slower for *A. lyrata*, possibly due to temperature-related reagent degradation or unknown contaminants in the DNA extraction. Over ~90h sequencing, 25,839 1D reads were generated with a total yield of 62.2Mbp; this included three days of sequencing at RBG Kew following a 16h drive, during which reagents and flowcell were stored sub-optimally (near room-temperature). BLASTN 2.4.0^29^ was used to remove 5,130 reads with identity to phage lambda. Data are given in Extended Data Table 2. The following week in a laboratory, NEBNext Ultra II sequencing libraries were prepared for four field-extracted samples (two individuals from each species) and sequenced on an Illumina MiSeq (300bp, paired end). In total, 11.3Gbp and 37.8M reads were generated (each ~ 8M reads and 2Gbp; see Supplementary Note 1).

### Field offline basecalling and bioinformatics in real-time

Offline basecalling using nanocall 0.6.13^30^ was applied to the R7.3 data as no offline R9 basecaller was available at the time. Basecalled reads were compared to the reference genomes of *A. thaliana* (TAIR10 release) and *A. lyrata subsp petraea* (1.0 release). In total, 119 reads were processed in real-time with six reads making significant hits by BLASTN that scored correctly: incorrectly for species ID in a 2:1 ratio. After the sequencer had been halted a larger dataset of 1,813 reads gave 281 hits, with correct: incorrect: tied identifications in a 223:30:28 ratio.

### Accuracy and mapping rates of short-and long-read data

Both lab-sequenced NGS reads (trimmed with Trimmomatic^31^) and untrimmed, field-sequenced RTnS reads were aligned to the appropriate reference genomes using the BWAv0.7.12-r1039^32^ and LASTv581^33^, to estimate depth of coverage and nominal error rate in mapped regions (see Supplementary Note 2). For all *A. thaliana* datasets (short and long-read), average mapped read depths were approximately equal to the gross coverage. MinION reads could be aligned to 53Mbp of the reference genome with LAST (approx. 50% of the total genome length). The nominal average error rate in these alignments was 20.9%). For both MinION and MiSeq datasets, mapping and alignment to the *A. lyrata* and *A. lyrata ssp. Petraea* assemblies was more problematic. For alignable MinlON reads, error rates were slightly higher than for *A. thaliana* at 22.5% and 23.5%, estimated against *A. lyrata* and *A. lyrata ssp. petraea* assemblies, respectively. We note that these assemblies are poorer quality than the *A. thaliana* TAIR10 release; total genome lengths differ (206Mbp and 202Mbp,) and contiguity is relatively poor in both (695 and 281,536 scaffolds).

### Determination of true-and false-positive detection rates, sensitivity, and specificity

Each of the four datasets (HTS and RTnS, for each species) was matched against two custom databases (the *A. thaliana* reference genome and the two draft *A. lyrata* genomes combined) separately with BLASTN, retaining only the best hit for each query. Queries matching only a single database were counted as positive matches for that species (Extended Data Table 4). Non-matching reads were treated as negative results (Supplementary Methods). Queries matching both databases were defined as positives based on: a) longest alignment length (L_T_); b) highest % sequence identities, c) longest alignment length counting only identities (L_I_), or c) lowest E-value. Test statistics for each of these metrics were simply calculated as the difference of scores (length ΔL_T_), *%* identities, identities (ΔL_I_), or E-value) between ‘true’ and ‘false’ hits. The statistical performance of these statistics (true-and false-positive rates, and accuracy) in putative analyses under varying threshold values were calculated and visualized using the ROCR package in R^34^. The high proportion of reads with significant hits to both species is expected given the close evolutionary relationships of the species. Analyses to determine the best statistics to discriminate between species using reads which aligned to both databases strongly indicated that difference in alignment lengths between the best discriminator, shown in Figure 2a-d and Extended Data Figures 2, 3 & 4. Overall these show that the difference in alignment length is a powerful indicator for both short-and long-read data at any threshold ≥ ~100bp. Furthermore, and surprisingly, at this and more conservative (greater difference) threshold, long-read field-sequenced reads had substantially more accuracy in true-and false-positive discrimination than short-read data. This suggests that this method provides a powerful means of species identification and we posit that the extremely long length of ‘true positive’ alignments compared with the natural length ceiling on false-positive alignments is largely responsible for this property.

### Accumulation curves for simulated identification

33,806 pairwise BLASTN hits obtained above in identification against *A. thaliana* and *A.lyrata* genomic reference databases were subsampled without replacement to simulate incremental accumulation of BLASTN hit data during progress of a hypothetical sequencing experiment producing 10,000 reads produced in total. 1,000 replicates were used to calculate means and variances for data accumulation in 0.1 log-increments from *r*=1 read to 10^4^ reads total. For each read, ΔL_I_, ‘number of identities bias’, was calculated as the difference (number of identities in *A. thaliana* alignment – number of identities in *A. lyrata* alignment). Each read was assigned to *A. thaliana* or not if it ΔL_I_ exceeded a given threshold, repeated at four possible values, L_threShold_ ={0, 1, 10, 100}. Mean and aggregate (total) ΔL_I_ values were also calculated for each replicate over the progress of the simulated data collection. Results are shown in Figure 3.

### *De novo* genome assembly

Short-read HTS data was assembled *de novo* using ABYSS v1.9.0^35^. A hybrid assembly with both HTS and RTnS datasets was performed with HybridSPAdes v3.5.0^36^. Assemblies were completed for *A. thaliana* (sample AT2a) and *A. lyrata* (sample AL1a). Assembly statistics were calculated in Quast v4.3^37^. Completeness of the final hybrid assemblies was assessed using CEGMA v2.5^38^. Results of *de novo* genome assemblies are given in Extended Data Table 5. Analyses of genome contiguity and correctness and conserved coding loci completeness indicated that assembly of HTS data performed as expected (20x coverage produced ~25,000 contigs covering approximately 82% of the reference genome at an N50 of 7,853bp). By contrast, the hybrid assembly of *A. thaliana* illumina MiSeq and Oxford Nanopore MinlON data significantly improved on the MiSeq-only assembly: 24,999 contigs reduced to 10,644; total assembly length increased to close to the length of the reference genome (119.0Mbp) with nearly 89% mappable; N50 and longest contig statistics both improved (N50 7,853 → 48,730bp) indicating better contiguity from the addition of long reads. Completeness of coding loci as estimated by CEGMA (Extended Data Table 5) greatly increased to ~99%. Long reads did not compromise the accuracy of high-coverage short-read data; basewise error rates were not significantly worse.

### Direct gene annotation of single unprocessed field-sequenced reads

The length of typical individual RTnS reads is of similar magnitude to genomic coding sequences. Consequently, useful phylogenomic information could potentially be obtained by annotating reads directly, without a computationally expensive genome assembly step. Raw, unprocessed *A. thaliana* reads were individually annotated directly without assembly via SNAP^39^. To verify which gene predictions were genuine, the DNA sequences (and 1kb flanking regions, where available) were matched to available *A. thaliana* (TAIR10) genes with default parameters. BLAST hits were further pruned based on quality (based on 1^st^-quartile quality scores: alignments length bias ΔL_T_ ≥ +570bp / *%* identities bias ≥ +78.68 / E-value bias ≥ 0), reducing the number of hits from 18,098 to 10,615. Sample read alignments and details of SNAP output BLAST score summary statistics are given in Supplementary Table 1 and encounter curves-through-time are shown in Figure 2e.

### Phylogenomics of raw-read-annotated *A. thaliana* genes

Predicted *A. thaliana* gene sequences were combined with a published phylogenomic dataset spanning 852 orthologous, single-copy genes in plants and algae^28^, downsampled to 6 representative taxa for speed: *Equisetum diffusum, Juniperus scopulorum, Oryza sativa, Zea mays, Vitis vinifera* and *A. thaliana.* Our putative gene models were assigned identity based on reciprocal best-hit BLASTN matching with the *A. thaliana* sequences in these alignments, yielding 207 matches, of which the top 56 were used for phylogenomic analysis (Supplementary Table 1), only 18 having no missing taxa in the Wickett *et al*.^28^ dataset. Alignments were refined using MUSCLE v3.8.31^40^ and trimmed with a 50% missing-data filter (using trimAL v1.4rev15^41^) then used to infer species trees in two ways: (i) single gene phylogenies inferred separately (using RAxML v7.2.8^42^) under the GTRCAT substitution model with 10 discrete starting trees then combined into a summary tree using TreeAnnotator v.1.7.4^43^; (ii) a species tree inferred directly from the data under the multispecies coalescent^44^, implemented in ^*^BEAST v2.4.4^45^ (with adequate MCMC performance confirmed using Tracer v1.5). A maximum clade credibility (MCC) tree was produced using TreeAnnotator v.1.7.4. Phylogenies inferred by orthodox (RAxML) and multispecies coalescent (^*^BEAST) methods are shown in Extended Data Figure 5 and agreed with each other and the established phylogeny presented in Wickett *et al.*^28^

## Extended Data

**Extended Data Figure 1.**
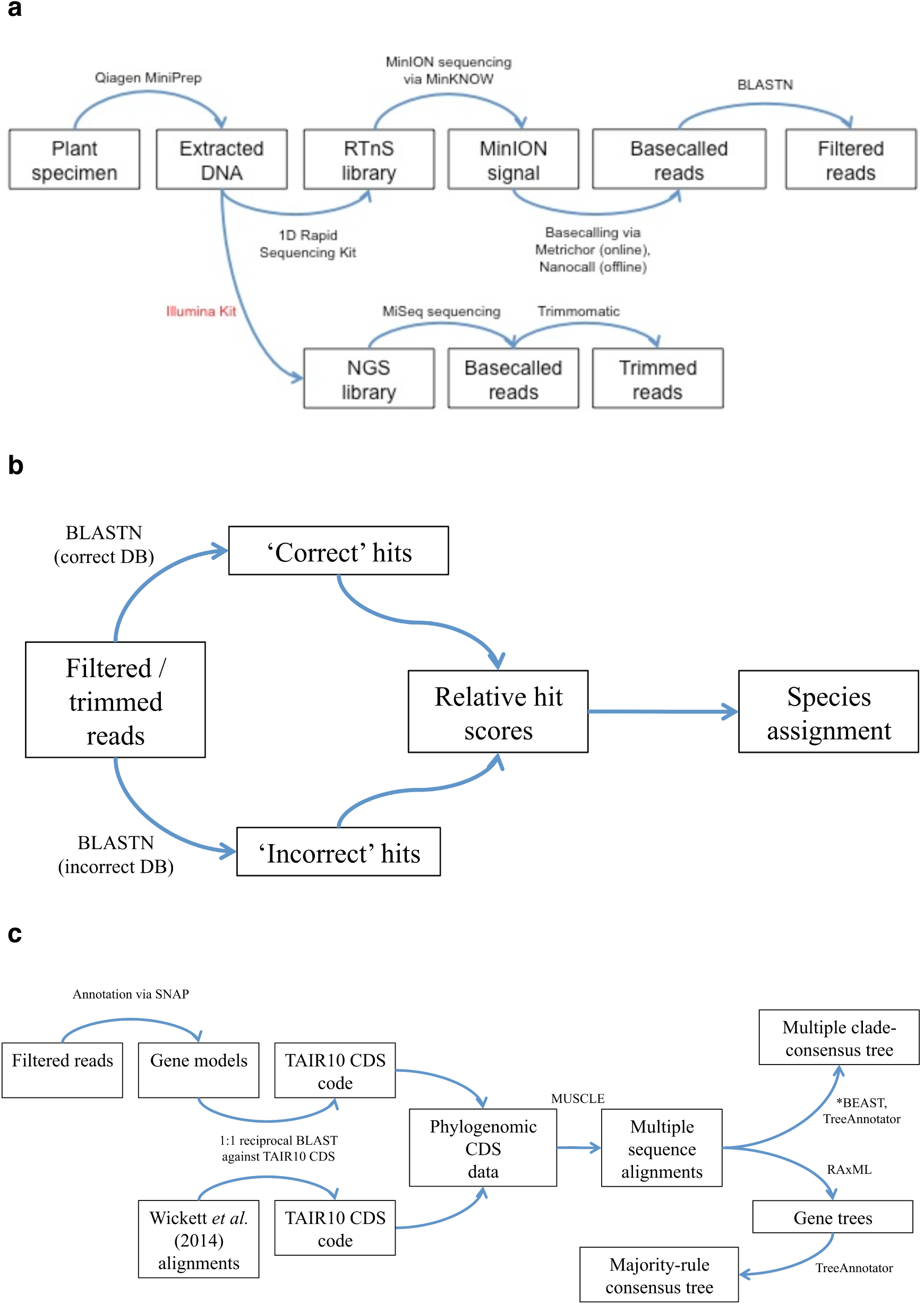
Schematic of experimental workflows. **a,** Sampling-to-sequencing workflow. **b,** Sample identification workflow via BLASTN. **c,** Outline for direct annotation of raw RTnS reads followed by phylogenomic inference. See Methods for details.

**Extended Data Figure 2.**
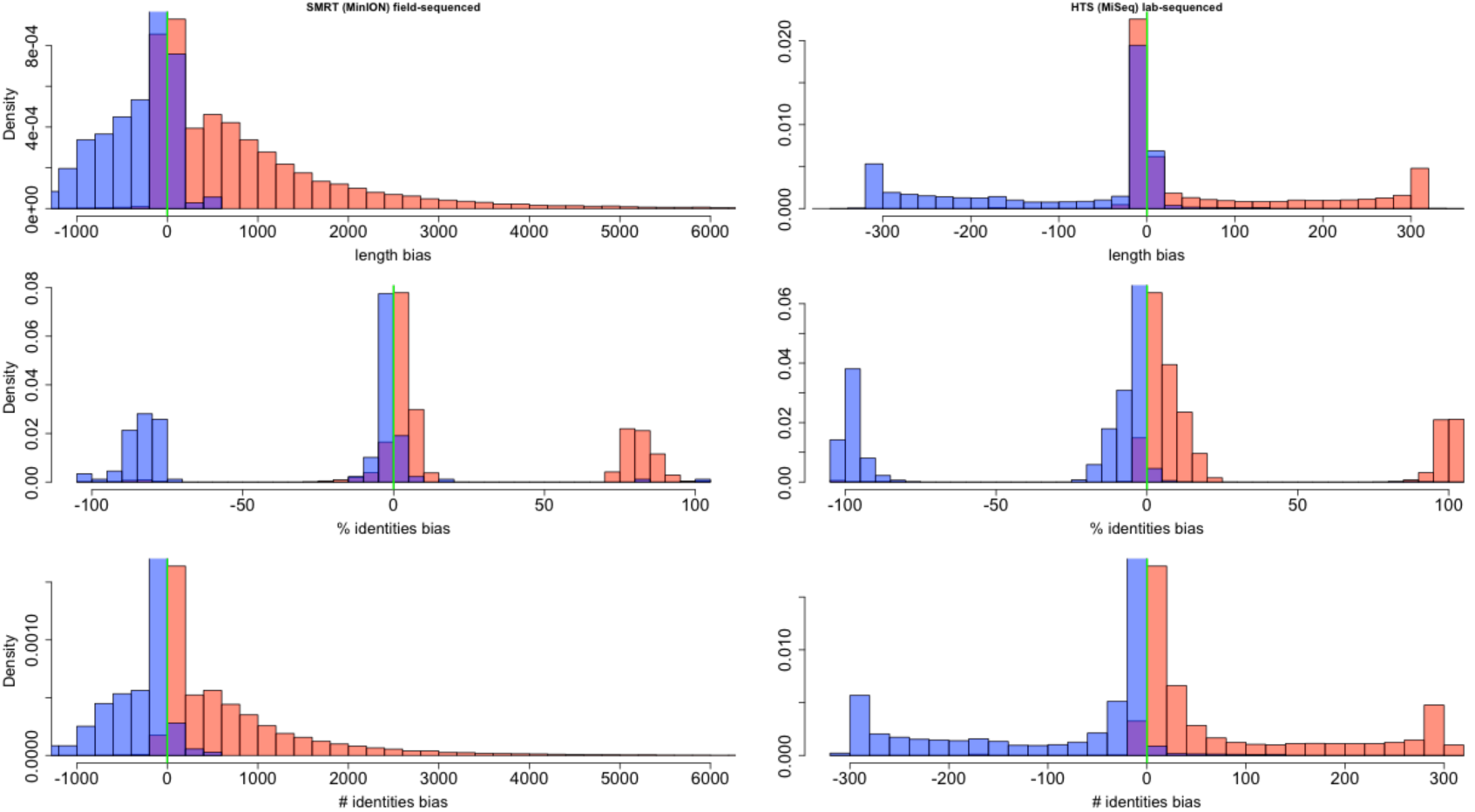
Distribution of difference statistics in BLASTN comparisons for species ID. Empirical distribution of test statistics for each-way congeneric sample ID (binary classification) using BLASTN evaluated for RTnS MinION (*left column*) and NGS MiSeq (*right column*) platforms. Difference (test) statistics were calculated for each alignment as (true positive (TP) score – false positive (FP) score) for each of: alignment length; % identities; and number of identities. Reads sequenced from *A. thaliana* samples (comprising nominal true positives and false-positives (contaminants) are shown in red; reads from *A. lyrata* samples (nominal true negatives) shown in blue. ‘True’ and ‘false’ distributions’ overlap is small, while alignment length and number of identities’ distributions are both unimodal, showing a simple cutoff-based classifier should perform well to discriminate between ‘true’ and ‘false’ cases.

**Extended Data Figure 3.**
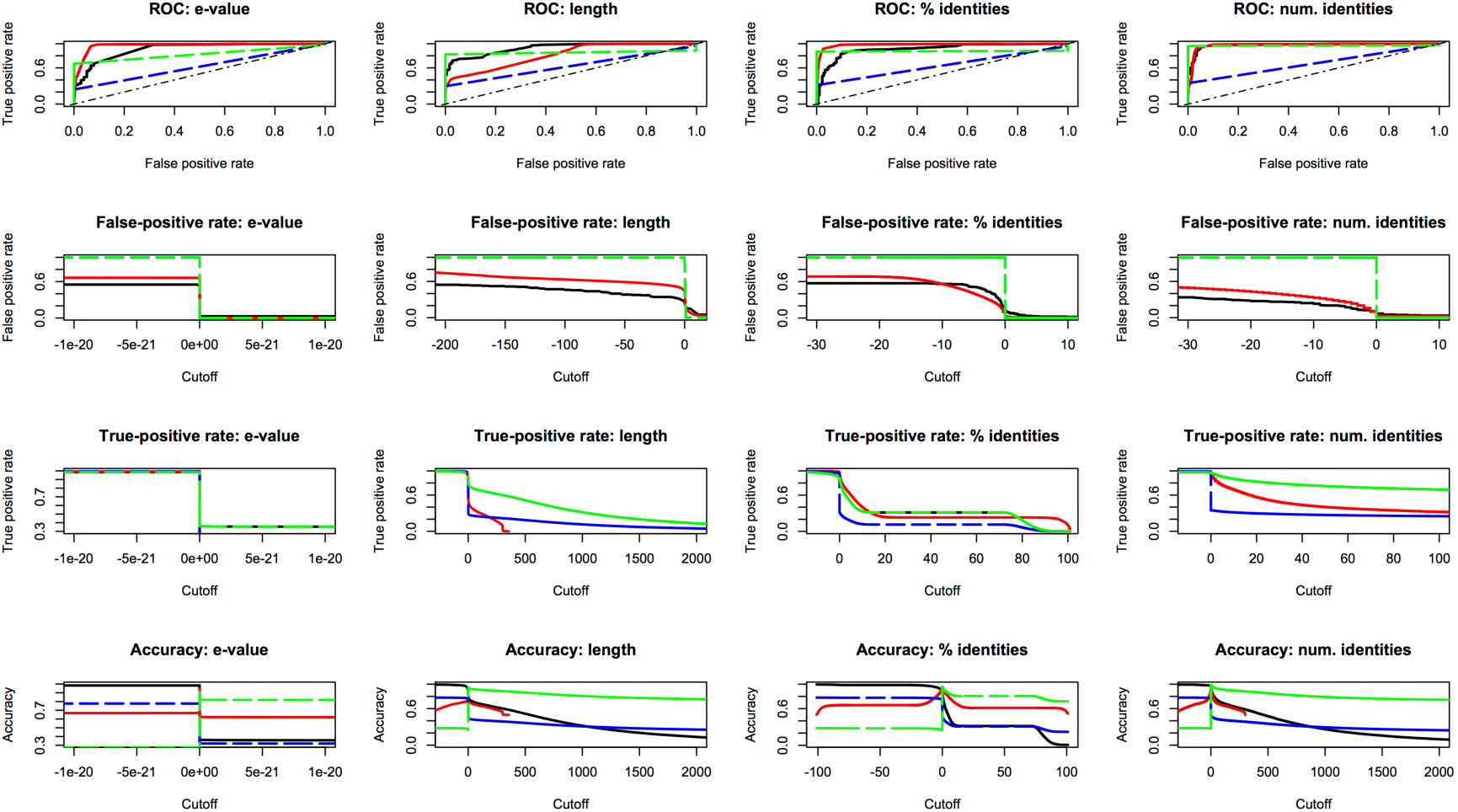
Performance of difference statistics in BLASTN comparisons for species ID. Performance of test statistics for each-way congeneric sample ID (binary classification) using BLASTN evaluated for MinION (*black*) and MiSeq (*red*) platforms. Difference (test) statistics were calculated amongst reads matching both databases for each alignment as (true positive (TP) score – false positive (FP) score) for each of e-value, length, % identities and number of identities. Reads matching only one or neither database were additionally included with either ‘all-false’ encoding (dashed green line) or mixed false and true encoding depending on sample origin (blue line; see Methods and Supplementary Information for details). RTnS reads’ true-and false-positive rates are comparable to, and in some cases better than, NGS reads’ performance; while the longer length of RTnS reads permits the use of high thresholds where greater confidence is desired (perhaps in the case of very closely related specimens). *Top row:* true-positive (TP) vs false-positive (FP) rate; classical receiver operating curve. *Second row:* FP rate with varying test statistic threshold. *Third row:* TP rate with varying test statistic threshold. *Bottom row:* Accuracy with varying test statistic threshold. Accuracy estimated as (TP+TN / (P + N)). *Columns (L-R):* Difference statistics for *e*-value, total alignment length, % identities, and number of identities, respectively.

**Extended Data Figure 4.**
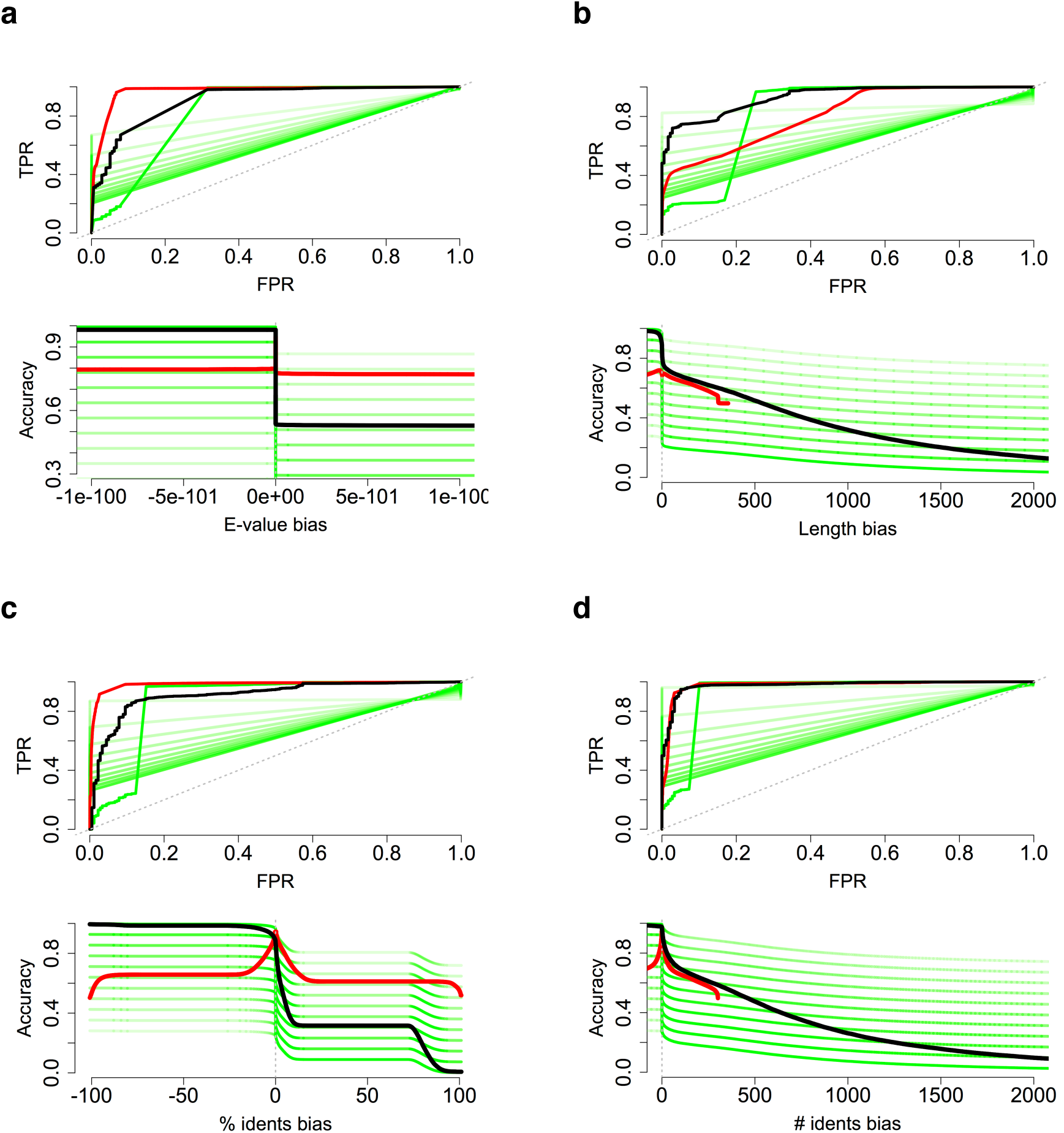
Modelling potential effect of incorrectly estimated TN/FN proportions. Red and black lines show empirically estimated statistical performance for species ID via BLASTN comparison of HTS and RTnS reads respectively (as for Extended Data Figure 3; see Methods for details). Reads that produced no hits to either database might represent false negatives (sequencing error, or genomic regions not represented in the reference genome BLAST databases) or true negatives (sequencing contaminants and sequencer noise). These nonmatching reads were to reflect ‘true negative’:’false negative’ (‘TN:FN’) mixtures in 10% increments shown from light to dark green shading. Plots (**a-d**) show results for bias statistics in *E-* value; total alignment length; % identities; and total alignment identities, respectively. Extreme TN:FN mixtures still display adequate true-positive vs. false-positive rates; empirical data is approximated by the 30% TN:FN mixture, approximately reflecting the proportions of *A. lyrata* to *A. thaliana* nonmatching reads in the dataset.

**Extended Data Figure 5.**
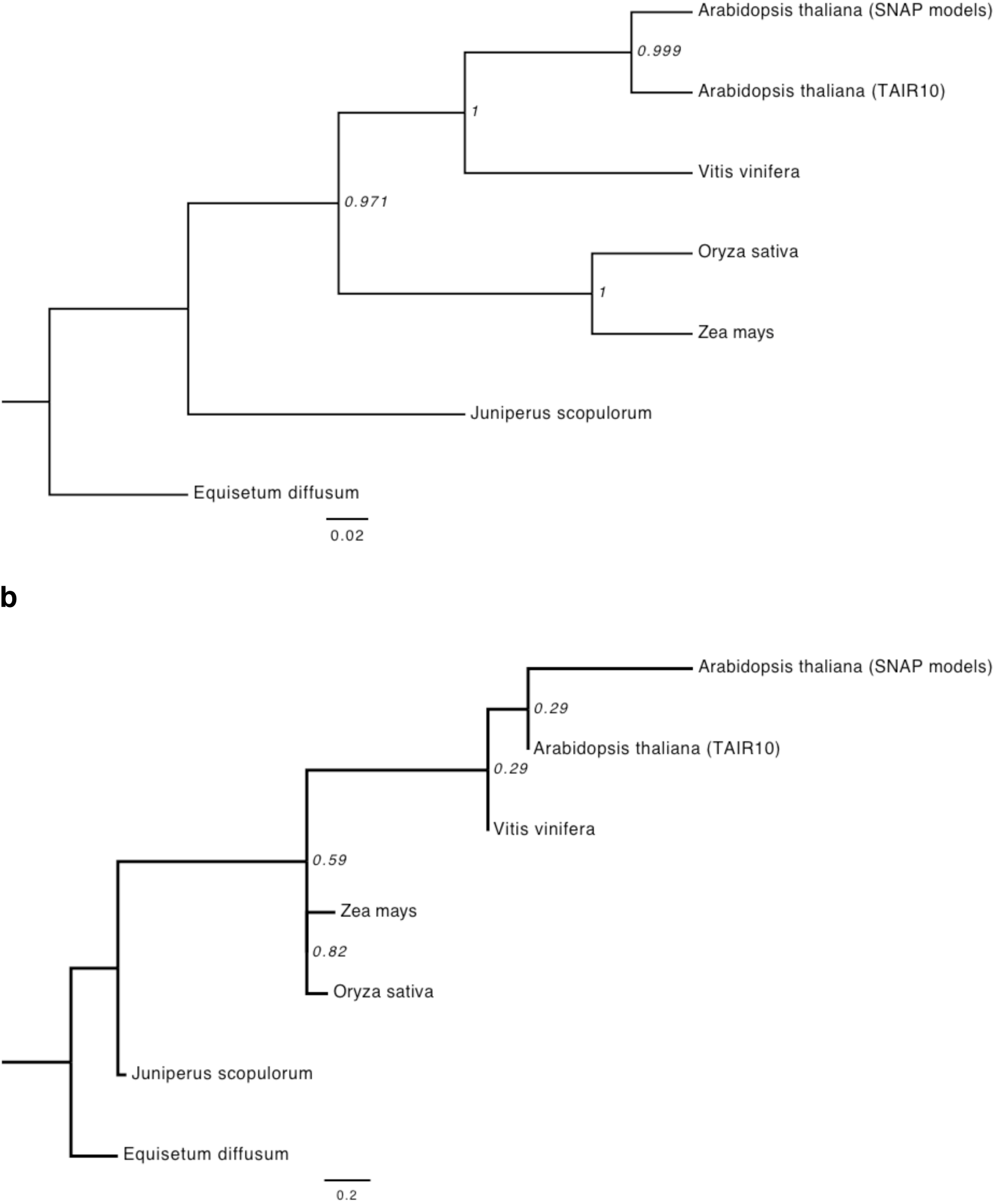
Phylogeny of species spanning major plant groups. Putative coding sequences recovered from single, unassembled raw RTnS reads using SNAP *ab initio* gene prediction could be aligned to existing phylogenomic data from other taxa and used to infer a phylogeny consistent with accepted plant relationships. **a,** Multispecies coalescent species tree inferred from 18 gene trees (genes predicted directly from raw nanopore reads). Inferred using multispecies coalescent implemented in BEAST 2.4.4; **b,** consensus species tree inferred by majority-rule from 18 gene trees, inferred with RAxML 7.2.8.

**Extended Data Table 1.** List of field-sequencing equipment.

| Item / model | Quantity | Supplier |
| --- | --- | --- |
| Laptops, Portege R830-1DZ | 2 | Toshiba |
| Laptop, MacBook Pro | 1 | Apple |
| Portable firewire HDD, 1Tb | 2 | LaCie |
| Portable AC generator, IMPAX IM800I 700W | 1 | ScrewFix |
| Thermal control textiles (socks) for MinION | Pair | ScrewFix |
| AC extension plugs, 4-way, 3-20M | 3 | Argos |
| Portable folding tables | 2 | Argos |
| Weather meter, Kestrel 5500 | 1 | KestrelMeters |
| Fluorometer, Quantus | 1 | Promega |
| Water bath, GD100 | 1 | Grant |
| Microcentrifuge, 5415C | 1 | Eppendorf |
| Glass thermometers, 300mm, range 263.3-383.3°K | 2 |  |
| Arduino Uno | 1 | Maplins |
| Thermal transducers, LM335AZ | 4 | RS Components |
| Polystyrene thermal control boxes, various sizes | 3 |  |
| Freezer coolpacks | 14 |  |
| Pipettes, Gilson: | - |  |
| P2 | 1 | ThermoFischer |
| P20 | 1 | ThermoFischer |
| P200 | 1 | ThermoFischer |
| P1000 | 1 | ThermoFischer |
| Waste containers | 2 |  |
| Laboratory consumables: | - |  |
| PCR tubes, thin-walled | Box | ThermoFischer |
| Eppendorf tubes, 1.5ml DNA lo-bind | Box | ThermoFischer |
| Eppendorf tubes, 2.0ml DNA lo-bind | Box | ThermoFischer |
| Plastic pestles | 20 |  |
| Sterilized sand | 5g |  |
| Reagent tubes, screw-top, 50ml | 10 | ThermoFischer |

**Extended Data Table 2.**
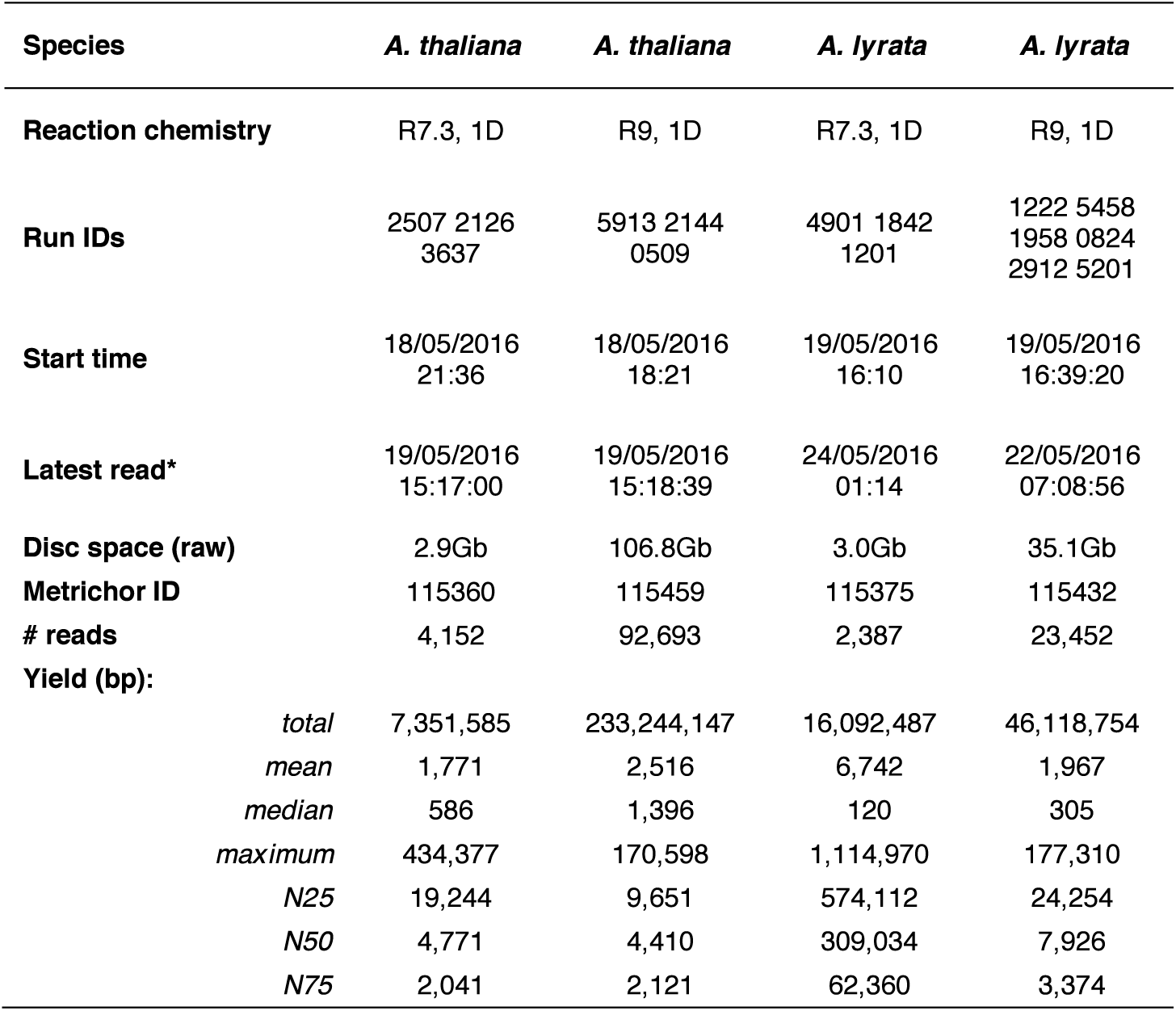
Performance of MinION sequencing runs. Data produced with field-extracted and field-sequenced in conditions ranging from 6-14°C and up to 100% humidity, producing genome-scale sequence data despite several pauses in sequencing to dismantle, relocate, and reassemble equipment. The two latest sequencing runs (*A. iyrata* samples) performed markedly worse than the two earliest runs (*A. thaliana*), possibly due to the impact of storage temperature (passively controlled, and steadily rising over the week) on reagent performance. Yield summary statistics refer to untrimmed raw reads, including phage-lambda experimental control in the case of *A. thaliana* R9 data (filtered from subsequent steps). See Methods for details. Note: ^*^Final *A. lyrata* sequencing phase performed in laboratory owing to time constraints on-site.

| Species | *A. thaliana* | *A. thaliana* | *A. lyrata* | *A. lyrata* |
| --- | --- | --- | --- | --- |
| Reaction chemistry | R7.3, 1D | R9, 1D | R7.3, 1D | R9, 1D |
| Run IDs | 2507 2126<br>3637 | 5913 2144<br>0509 | 4901 1842<br>1201 | 1222 5458<br>1958 0824<br>2912 5201 |
| Start time | 18/05/2016<br>21:36 | 18/05/2016<br>18:21 | 19/05/2016<br>16:10 | 19/05/2016<br>16:39:20 |
| Latest read* | 19/05/2016<br>15:17:00 | 19/05/2016<br>15:18:39 | 24/05/2016<br>01:14 | 22/05/2016<br>07:08:56 |
| Disc space (raw) | 2.9Gb | 106.8Gb | 3.0Gb | 35.1Gb |
| Metrichor ID | 115360 | 115459 | 115375 | 115432 |
| # reads | 4,152 | 92,693 | 2,387 | 23,452 |
| Yield (bp): |  |  |  |  |
| total | 7,351,585 | 233,244,147 | 16,092,487 | 46,118,754 |
| mean | 1,771 | 2,516 | 6,742 | 1,967 |
| median | 586 | 1,396 | 120 | 305 |
| maximum | 434,377 | 170,598 | 1,114,970 | 177,310 |
| N25 | 19,244 | 9,651 | 574,112 | 24,254 |
| N50 | 4,771 | 4,410 | 309,034 | 7,926 |
| N75 | 2,041 | 2,121 | 62,360 | 3,374 |

**Extended Data Table 3.**
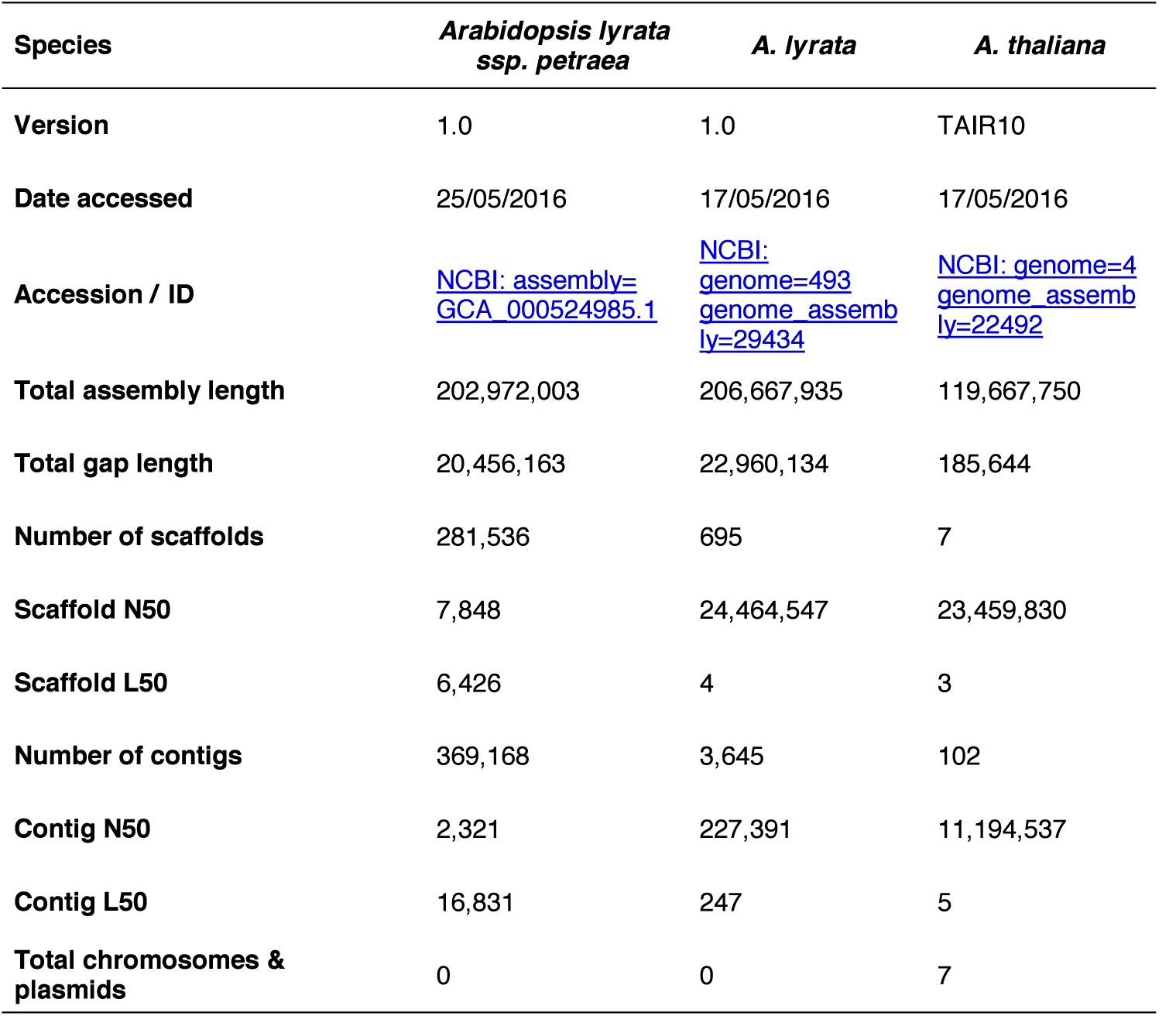
Statistics of reference genomes used. *A. thaliana* TAIR10 release is considerably more complete than either of the draft *A. iyrata* assemblies.

| Species | *Arabidopsis lyrata*<br>*ssp. petraea* | *A. lyrata* | *A. thaliana* |
| --- | --- | --- | --- |
| Version | 1.0 | 1.0 | TAIR10 |
| Date accessed | 25/05/2016 | 17/05/2016 | 17/05/2016 |
| Accession / ID | NCBI: assembly=<br>GCA_000524985.1 | NCBI:<br>genome=493<br>genome_assemb<br>ly=29434 | NCBI: genome=4<br>genome_assemb<br>ly=22492 |
| Total assembly length | 202,972,003 | 206,667,935 | 119,667,750 |
| Total gap length | 20,456,163 | 22,960,134 | 185,644 |
| Number of scaffolds | 281,536 | 695 | 7 |
| Scaffold N50 | 7,848 | 24,464,547 | 23,459,830 |
| Scaffold L50 | 6,426 | 4 | 3 |
| Number of contigs | 369,168 | 3,645 | 102 |
| Contig N50 | 2,321 | 227,391 | 11,194,537 |
| Contig L50 | 16,831 | 247 | 5 |
| Total chromosomes & plasmids | 0 | 0 | 7 |

**Extended Data Table 4.** Sample identification via BLASTN. Individual RTnS and NGS reads were aligned to *A. thaliana* and *A. lyrata* databases (designated TRUE or FALSE depending on sample origin) with BLASTN, keeping the single best-hit alignment for each database. More ‘1-way’ (only one database matched) hits to TRUE than FALSE databases accumulated in all sample / technology combinations. Amongst ‘2-way’ hits, positive differences in the metrics were consistent with correct sample identification. For RTnS reads differences between TRUE and FALSE hits were considerably larger than amongst HTS reads (by an order of magnitude for length or number of identities), showing that confident identification could be made with fewer RTnS reads. Notes: ^*^*A. lyrata* and *A. lyrata ssp. petraea* databases combined, see Methods; †Total number of reads matching only conspecific database (‘true-positives’); ‡Total number of reads matching only pairwise-compared database (‘false-positives’ in the case of a mixed /multiplexed sample, or ‘false-negatives’ in the case of a single sample); §Total number of reads matching both databases; ¶Total number of reads with no hits in either comparison, e.g. ‘false-negatives’; #Difference statistics for each query read calculated as (score conspecific comparison – score congener comparison), for BLASTN alignment length, alignment identities, alignment % identities and *E*-value; ⋆Mean bias across all reads.

| Sample | *A. thaliana* | *A. lyrata* | *A. thaliana* | *A. lyrata* |
| --- | --- | --- | --- | --- |
| **“TRUE” database** | *A. thaliana* | *A. lyrata*<br>combined* | *A. thaliana* | *A. lyrata*<br>combined* |
| **“FALSE” database** | *A. lyrata*<br>combined* | *A. thaliana* | *A. lyrata*<br>combined* | *A. thaliana* |
| **Source data** | ONT 1D | ONT 1D | MiSeq | MiSeq |
| **# Reads, total** | 91,715 | 25,839 | 9,476,598 | 9,659,489 |
| **# Reads with BLASTN hits:** |  |  |  |  |
| 1-way TRUE<sup>†</sup> | 10,322 | 76 | 2,140,403 | 2,907,921 |
| 1-way FALSE<sup>‡</sup> | 378 | 2 | 53,056 | 24,329 |
| 2-way BOTH<sup>§</sup> | 22,386 | 101 | 7,098,032 | 6,256,969 |
| 0-way ZERO<sup>¶</sup> | 58,629 | 25,660 | 185,107 | 470,270 |
| **Proportion of reads:** |  |  |  |  |
| 1-way TRUE | 0.113 | 0.003 | 0.226 | 0.301 |
| 1-way FALSE | 0.004 | 0.000 | 0.006 | 0.003 |
| 2-way BOTH | 0.244 | 0.004 | 0.749 | 0.648 |
| 0-way ZERO | 0.639 | 0.993 | 0.020 | 0.049 |
| **Biases<sup>#</sup>:** |  |  |  |  |
| Mean length | 1,323.87 | 698.25 | 83.61 | 108.99 |
| Mean identities | 1,115 | 575 | 96 | 117 |
| Mean % identities | 37.97 | 61.41 | 30.16 | 42.38 |
| Mean E-values | 4.80E-07 | 1.05E-04 | 1.07E-08 | 3.52E-09 |

**Extended Data Table 5.** Performance of *de novo* genome assembly. Field-extracted DNA material was of sufficient quality to enable a *de novo* assembly with lab-sequenced NGS data. Furthermore, field-sequenced RTnS reads considerably augmented the NGS data in hybrid assembly, greatly improving contiguity and estimated coding loci coverage substantially without substantially raising basewise error rates. Notes: *^*^de novo* genome assemblies used either lab-sequenced short-read HTS data only (Abyss) or both HTS and field-sequenced RTnS datasets (Hybrid-SPAdes). †TAIR10 release. ‡INSDC: *A. lyrata*: ADBK00000000.1 (Hu *et al.*, 2011); *A. lyrata ssp. petraea:* BASP00000000.1 (Akama *et al.*, 2014). §Assembly statistics calculated using QUAST 4.0. ⋆Approximate completeness of coding loci assessed via CEGMA. See Methods for details.

| Species | *Arabidopsis thaliana* |  | *A. lyrata ssp. petraea* |  |
| --- | --- | --- | --- | --- |
| Data | MiSeq | MiSeq + MinION | MiSeq | MiSeq + MinION |
| Assembler * | Abyss | hybridSPAdes | Abyss | hybridSPAdes |
| Illumina MiSeq NGS reads, 300bp paired-end | 8,033,488 | 8,033,488 | 8,143,010 | 8,143,010 |
| NGS total yield | 2,418,079,888 | 2,418,079,888 | 2,451,046,010 | 2,451,046,010 |
| Oxford Nanopore MinION RTnS reads, R7.3 + R9, N50 ~ 4,410bp | n/a | 96,845 | n/a | 25,839 |
| RTnS reads total yield | n/a | 240,597,532 | n/a | 62,211,241 |
| # contigs | 24,999 | 10,644 | 37,568 | 85,599 |
| Largest contig | 89,717 | 413,462 | 101,114 | 38,313 |
| Total length | 106,455,313 | 119,031,857 | 151,562,895 | 117,256,694 |
| Reference length | 119,667,750 <sup>†</sup> | 119,667,750 | 183,707,801<sup>‡</sup> | 183,707,801 |
| GC content (%) | 35.97 | 36.20 | 36.16 | 36.55 |
| N50 <sup>§</sup> | 7,853 | 48,730 | 9,605 | 1,686 |
| Unaligned length | 7,121,882 | 6,737,059 | 36,669,847 | 35,287,390 |
| Genome fraction (%) | 82.0 | 88.7 | 53.4 | 43.7 |
| Duplication ratio | 1.01 | 1.058 | 1.17 | 1.02 |
| # N's per 100 kbp | 1.72 | 5.41 | 0.22 | 7.09 |
| # mismatches / 100 kbp | 518 | 588 | 1,297 | 1,097 |
| # indels / 100 kbp | 120 | 130 | 334 | 271 |
| Largest alignment | 76,935 | 264,039 | 44,515 | 17,201 |
| Total aligned length | 98,382,255 | 108,086,256 | 100,502,092 | 80,814,492 |
| *Coding loci completeness<sup>¶</sup>:* |  |  |  |  |
| # genes, 'complete' | **219** | **245** | n/a | n/a |
| % genes, 'complete' | 88.31% | 98.79% |  |  |
| # genes 'partial' | **238** | **246** | n/a | n/a |
| % genes, 'partial' | 95.97% | 99.19% |  |  |

